# A novel cell-free method to culture *Schistosoma mansoni* from cercariae to juvenile worm stages for *in vitro* drug testing

**DOI:** 10.1101/341669

**Authors:** Sören Frahm, Anisuzzaman, Fabien Prodjinotho, Admar Verschoor, Clarissa Prazeres da Costa

## Abstract

**Background:** The arsenal in anthelminthic treatment against schistosomiasis is limited and relies almost exclusively on a single drug, praziquantel (PZQ). Thus, resistance to PZQ could constitute a major threat. Even though PZQ is potent in killing adult worms, it has been shown to be limited in its activity against earlier developmental stages. Current *in vitro* screening strategies for new drugs depend on newly transformed schistosomulae (NTS) for initial hit identification, thereby limiting sensitivity to new compounds predominantly active in later developmental stages. Therefore, the aim of this study was to establish a highly standardized, straightforward and reliable culture method to generate and maintain advanced larval stages *in vitro*. We present here how this method can be a valuable tool to test drug efficacy at each discrete intermediate larval stage, reducing the reliance on animal use (3Rs).

**Methodology/principal findings:** Cercariae were mechanically transformed into skin-stage (SkS) schistosomulae and successfully cultured under serum-free and cell-independent conditions for up to four weeks with no loss in viability. Under these conditions, larval development halted at the lung-stage (LuS). Addition of human serum (HSe) propelled further development into juvenile worms within eight weeks. Skin and lung stages, as well as juvenile worms, were submitted to 96-well format drug screening assays using known anti-schistosomal compounds such as PZQ, oxamniquine (OXM), mefloquine (MFQ) and artemether (ART). Our findings showed stage-dependent differences in larval susceptibility.

**Conclusion:** With this robust and highly standardized *in vitro* assay, important developmental stages of *S. mansoni* up to juvenile worms can be generated and maintained over prolonged periods of time. The phenotype of juvenile worms when exposed to reference drugs was comparable to previously published works for *ex vivo* harvested adult worms. Therefore, this *in vitro* assay can help reduce reliance on animal experiments in the search for new anti-schistosomal drugs.

**Author Summary:** Schistosomiasis remains a major health threat, predominantly in developing countries. Even though there has been some progress in search of new drugs, praziquantel remains the only available drug. Probably the most important advance in the search for new drugs was *in vitro* transformation of cercariae and their subsequent culture. However, hit identification in compound screenings is exclusively tested in skin stage parasites and is only confirmed for more mature worms in a subsequent step. This is in part due to the lack of an easy culture system for advanced-stage parasites. We present here a reliable and highly standardized way to generate juvenile worms *in vitro* in a cell-free culture system. The inclusion of *in vitro* drug tests on advanced-stage parasites in initial hit identification will help to identify compounds that might otherwise be overlooked. Furthermore, the ability to continuously observe the parasite’s development *in vitro* will provide an important platform for a better understanding of its maturation in the human host. Taken together, this opens up new avenues to investigate the influence of specific cell types or host proteins on the development of *Schistosoma mansoni* and provides an additional tool to reduce animal use in future drug discovery efforts (3Rs).

## Introduction

Schistosomiasis, a chronic and debilitating helminthic disease, is one of the most important neglected tropical diseases (NTD). The WHO estimates that currently more than 206 million people are infected and in need of preventive chemotherapy world-wide [1] and over 200,000 people die each year due to the sequelae of the disease [2, 3]. Among the parasitic diseases, schistosomiasis is often considered only second in importance to malaria [2] and thus a major public health menace. Therefore, the WHO aims to eliminate schistosomiasis as a public health problem globally by 2025 [4, 5]. Implementation of safe water, sanitation and hygiene (WASH) strategies [6], intensified case management, veterinary public health, vector control and mass drug administrations (MDAs) are all crucial to reduce the disease burden [5]. Of all these approaches, MDA dominates national control programs thanks to the excellent safety and efficacy profile of praziquantel (PZQ), the only currently available drug [7, 8], as well as its low cost per treated individual [9]. However, the reliance on PZQ also raises concerns about emerging resistance should the drug pressure increase. Resistance against PZQ has already been observed in experimental models [ 10] while a decrease in drug efficacy has been observed in the field [7, 11, 12]. To be prepared for the emergence of resistant strains of schistosomes and to support the elimination of schistosomiasis, new drugs and complementary strategies, such as vaccines, are of imminent importance. To identify new drugs and new vaccine targets, *in vitro* assays of larval and adult worm stages are paramount for high throughput testing and simultaneously for reducing reliance on *in vivo* or *ex vivo* experiments in accordance with the 3Rs (replacement, reduction and refinement) of animal testing [13].

The current cultivation protocols for *in vitro* generated larval stages such as the schistosomulae rely on the supplementation of fetal calf serum (FCS) for short term culture [14, 15] or on the supplementation with FCS, human serum (HSe), erythrocytes and peripheral blood mononuclear cells (PBMCs) for long-term culture and *in vitro* juvenile worm development [16]. Such non-standardized culture conditions are prone to variability of serum batches, rely on the continuous supply of fresh human blood and make it difficult to isolate pure schistosomula-derived soluble antigens during *in vitro* culture or to investigate the role of specific serum proteins in the development of the parasite. In addition to these hindrances and to avoid inter-assay or inter-laboratory fluctuations, well-defined and standardized culture conditions independent of serum and cell supplementations are needed. Moreover, simplifying culture conditions as well as the generation and handling of advanced-stage parasites opens new possibilities in the search for new drugs and facilitates the upscaling of existing drug screening strategies.

To establish a robust *in vitro* assay, it is important to continuously imitate the parasites *in vivo* development. This development within the final host is quite complex and occurs over a period of seven weeks. After penetration of the skin by the cercariae, the schistosomulae can be found for approx. three days within the skin before they start migrating through the host’s vasculature. This skin stage (SkS) is followed by traversing the capillaries of the lung where the majority of the parasites can be found on day 7 after infection. To facilitate their migration, the parasites become longer, slenderer and more active. The lung stage (LuS) schistosomulae continue their journey to the portal and mesenteric veins following the bloodstream. There they initially undergo several morphological changes. The gut is formed, the parasites initiate feeding and they start growing. This early liver stage (LiS) is followed by the late LiS characterized by a drastic increase in length and prominent oral and ventral suckers. These juvenile worms (late LiS) then start to pair up and become fertile upon which the oviposition starts approximately 35 days after infection [17].

Treatment with PZQ, although initially with good efficacy, does not diminish the high reinfection rates encountered in the field [18]. This is partly due to the inability of PZQ to efficiently target early larval stages of the parasite [19, 20]. In the current search for new anti-schistosomal drugs, a two-step strategy is used. Firstly, SkS schistosomulae are tested immediately following their transformation from cercariae and then, once an active compound has been identified, it is tested mainly on *ex vivo* cultured adult worms that have been isolated from infected hamsters or mice [14, 21-23]. Subsequent larval stages like the LuS, early and late LiS are omitted as potential drug targets. Therefore, future compounds with an activity predominantly directed against juvenile and adult stages, such as PZQ, might be overlooked. Thus, a highly standardized and robust way to generate advanced larval stages of *Schistosoma mansoni* provides an opportunity to incorporate initial advanced-stage schistosomula testing into the current drug screening strategies to meet the desired compound target profile (TCP).

We disclose here a new serum- and cell-free cultivation method of newly transformed schistosomulae (NTS) up to the LuS and a cell-free cultivation method up to late LiS juvenile worms of *S. mansoni*. Our culture system allows the detection of stage-dependent differences in the activity of drugs with known anti-schistosomal properties. We used PZQ, the only currently available drug; Oxamniquine (OXM), the standard drug to treat schistosomiasis caused by *S. mansoni* prior to the advent of PZQ, and two antimalarial drugs that have recently been described to have anti-schistosomal properties, Mefloquine (MFQ) and Artemether (ART) [16, 24]. Therefore, the aim of this study was to establish a highly standardized, straightforward and reliable culture method as the basis for integrating drug screenings of advanced larval stages in initial hit identification. For verification of this cell-free cultivation assay, we tested compounds with already known anti-schistosomal properties against SkS schistosomulae. We hereby demonstrate that more advanced developmental stages such as LuS parasites and juvenile worms (LiS) can also be cultured and tested *in vitro* under cell-free conditions.

## Material and Methods

### NTS generation

Cercariae of a Brazilian strain of *S. mansoni* were harvested from infected *Biomphalaria glabrata* snails and used for mechanical transformation into NTS as described before [25]. Briefly, cercariae were incubated 30 min on ice then centrifuged at 1932 × g for 3 min at 4°C. The pellet was resuspended in Hank’s basal salt solution (HBSS) (Sigma-Aldrich, Germany) supplemented with 200 U/ml Penicillin and 200 µg/ml Streptomycin (Sigma-Aldrich) and transformed by mechanical stress applied by pipetting and vortexing, which was confirmed by microscopy (10x magnification). Separation of tails and cercarial bodies was accomplished by repeated sedimentation in ice-cold HBSS. The NTS were then transferred to the various culture media.

### NTS culture

NTS (100 NTS in 150 µl) were maintained in commercially available cell culture media such as HybridoMed Diff 1000 (HM) (Biochrom GmbH, Germany), Medium 199 (M199), Dulbecco’s Modified Eagle Medium (DMEM) or RPMI 1640 (Sigma-Aldrich, Germany) supplemented with 200 U/ml penicillin, and 200 µg/ml streptomycin (Sigma-Aldrich) in a 96- well flat bottom tissue culture plate (Corning Incorporated, USA) and incubated at 37°C in 5% CO_2_ and humidified air for up to 4 weeks. Medium was replaced every 7 days.

### Scoring criteria for the viability score

Viability of NTS was scored using an Axiovert10 microscope (Zeiss, Germany). The scoring system was adapted from the Swiss TPH [25] and the WHO-TDR [23]. The score represents the average of all NTS in a well. For the scoring, three main criteria were assessed: motility, morphology and granularity. The score was applied ranging from 0 (dead parasites, no movement, heavy granulation, blurred outline, rough outer tegument and blebs) to 1 (very reduced motility, rough outer tegument with some blebs) to 2 (reduced motility or increased uncoordinated activity, slight granularity, intact tegument with slight deformations) and finally 3 (regular smooth contractions, no blebs and a smooth outer surface, no granulation with clear view of internal structures). Viability was scored at indicated time points. Since the final score was as an average of all parasites in each well (n=3), the score points were applied in 0.25 steps (S1 Fig).

### Growth promotion with human serum

Blood sampling of HSe was prepared from blood of consenting healthy volunteers with no previous history of schistosomiasis upon written consent. Fresh blood was left at room temperature for 30 min to clot, then centrifuged at 1845 × g for 20 min and serum was collected and pooled from 6 individuals and stored at −20°C until further use. NTS were incubated (100 in 150 µl) in pure HM and DMEM as controls, or with HSe in different concentrations (1-50%) at 37°C for 8 weeks in a 96-well plate. Medium was changed weekly and viability was scored on day 1, 3 and 7 post transformation (p.t.) and then again, every 7 days. Developmental stages were determined via bright field microscopy using an inverted Axiovert 10 microscope (Zeiss).

### *In vitro* drug testing

PZQ (Merck, Germany), OXM, MFQ and ART (Sigma-Aldrich, Germany) were dissolved in DMSO depending on drug solubility (PZQ 10mg/ml, OXM 5 mg/ml, MFQ 33.3 mg/ml, ART 10 mg/ml) and stored at 4°C until use. NTS were cultured in HM supplemented with (SkS, LuS and LiS) or without (SkS and LuS) 20% HSe. To test the drug sensitivity of the distinct developmental stages, we incubated SkS (24-hour-old NTS), LuS (7-day-old NTS) and LiS (6- week-old NTS) parasites with PZQ, OXM, MFQ and ART at different concentrations (1, 10, 100 µg/ml). Before the addition of the drugs, a medium exchange was performed. Scoring was performed before (0 h) and 3, 24, 48, 72 and 168 h after treatment (a.t.).

### Statistics

For statistical analysis of the experiments to determine the optimal culture medium as well as the HSe concentration, a Krusal-Wallis test was performed on day 1 and 3, week 1, 4 and 8 p.t. and if significant (p ≤ 0.05), the data was further analyzed by employing Mann-Whitney U tests comparing pure medium with serum-supplemented medium followed by a Bonferroni correction. The reportable results are shown as a mean ± SD of the viability score points of three pooled independent experiments with three replicates each. Statistical testing was done with IBM SPSS Statistics 24 (IBM). For the creation of the graphs, PRISM 5 (GraphPad) was used.

For statistical analysis of the drug treatment experiments, Mann-Whitney U testing was performed to determine statistically significant differences between the DMSO control and the treated groups. The reportable results are shown as a mean ± SD of the viability score points of three pooled independent experiments with three replicates each. Statistical testing was done with IBM SPSS Statistics 24 (IBM). Graphs were generated using the Graphs PRISM 5 (GraphPad) was used.

## Results

### Serum- and cell-free medium ensures a high viability of *S. mansoni* NTS in long-term culture

To generate and maintain NTS under serum- and cell-free conditions, we cultivated NTS immediately following transformation in different highly standardized and commercially available culture media such as HM, DMEM, RPMI and M199 (Fig 1A, B) supplemented only with 200 U/ml penicillin and 200 µg/ml streptomycin. Over a period of four weeks, regular visual viability scoring of the parasites was performed following defined criteria adapted from previously published works [23] (Suppl. 1). In DMEM, NTS survived well for at least four weeks. On day 3 p.t., their viability peaked (2.6 ± 0.1), characterized by an increased motility and hardly any observable granularity. Then viability declined slightly (2.0 ± 0.0) due to the death of single NTS and slight internal granulation on day 7 after which the viability stabilized throughout the remainder of the experiment (Fig 1A, B). In HM, NTS initially scored (2.0 ± 0.3) on day 3 p.t., slightly lower compared to those in DMEM. However, from one week p.t. onwards, NTS in HM had already surpassed those kept in DMEM in the viability scoring (2.7 ± 0.3) due to the absence of granularity and increase of regular, steady movement. NTS in HM stayed viable throughout the experiment (four weeks) (Fig 1A, B). Compared to these two media (HM and DMEM), M199 and RPMI supported viability of NTS rather poorly. In M199, viability already started to drop (1.8 ± 0.3) on the third day p.t., characterized by the majority of the NTS being heavily damaged or already dead. By week 3 p.t., all NTS had died. (Fig 1A, B) In RPMI, the viability declined even faster with the majority of the NTS already heavily damaged or dead on day 3 p.t. and the death of all NTS by week 2 p.t. (Fig 1A, B). In all conditions, NTS were initially oval shaped on day 1 to day 3, resembling the SkS. Skin-stage schistosomulae were characterized by a stumpy looking oval outline with regular elongations and contractions either straight or to the side and were 105.7 ± 7.8µm in length (S3 Fig). By one week p.t., the surviving NTS had developed LuS characteristics. LuS NTS were characterized by a more elongated and slenderer form with an increased activity compared with the SkS. The parasites’ movement was still characterized by regular elongation and contraction in changing directions with an average length of 184.4 ± 45.5µm (Fig 1B and S3 Fig) [26]. In all media, however, development of surviving NTS halted after the first week of culture and, therefore, remained at the LuS stage for the remainder of the experiment. Dead NTS, in M199 or RPMI medium, displayed massive granulation internally and an irregular outer tegument. In contrast, healthy NTS, in HM and DMEM, were elongated and slender with a clear interior and a well contrasted outline (Fig 1B). Taking all of this into consideration, HM is the most suitable medium for long-term culture of LuS NTS without any serum or cell supplementation; however, development under these conditions is halted in the LuS.

**Fig 1.**
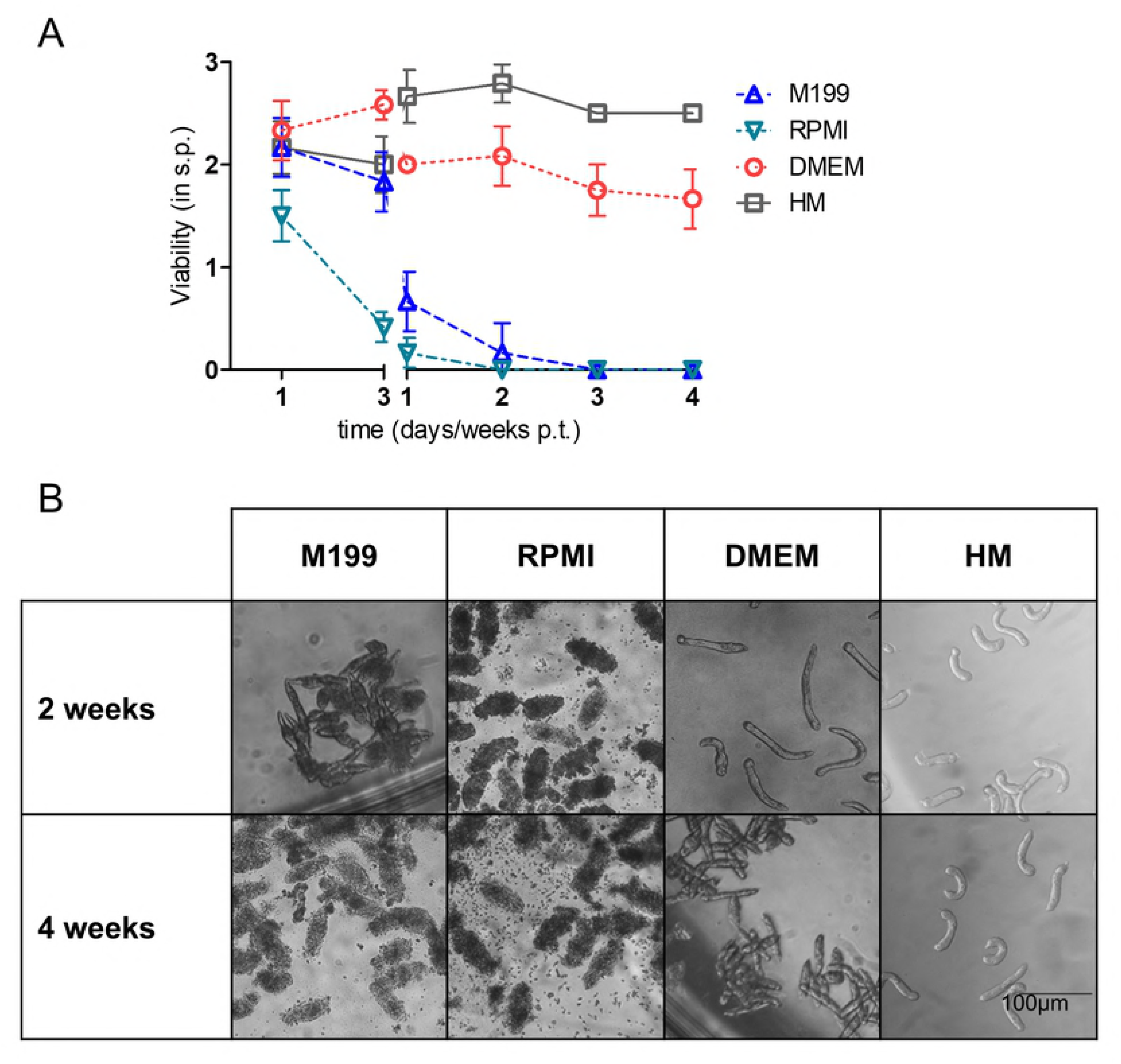
Culture media selectively ensure long-term viability of *S. mansoni* NTS without serum or cell supplementation. NTS were cultured in HM, M199, DMEM or RPMI for 4 weeks. (A) The viability of the parasites was scored at the indicated time points and (B) the morphology was observed under light microscopy. Photomicrographs were taken 2 and 4 weeks p.t. via 10x magnification. Results are representative of at least three independent experiments. Each data point is shown as mean ± SD of at least three biological replicates. p.t., post transformation; s.p., score points; HM, HybridoMed Diff 1000.

### HSe enhances NTS viability *in vitro* and induces development past the LuS up to juvenile worms

To test whether, in the absence of any cell supplementations, LuS NTS could develop further into LiS juvenile worms by exclusively adding HSe, the most important definite host [2, 27], we supplemented HM and DMEM with 20% HSe. In both serum-free controls, we observed a decline in viability after the fourth week of observation (Fig 2A). In serum-free HM and DMEM controls, only SkS and LuS schistosomulae were observed alongside some dead parasites. In HM and DMEM, all SkS that survived, developed to the LuS by the first week p.t. Until the fourth week p.t., around 49% (48.7 ± 4.2 dead out of 99.7 ± 10.6 total parasite count/well) of the NTS died in DMEM compared to only around 21% (39.0 ± 9.0 dead out of 185.7 ± 14.0 total) in HM. Following the fourth week p.t., a steady increase in dead NTS was observable in both conditions (Fig 2B, D). Addition of 20% HSe to the culture stopped the drop of viability and increase of dead NTS after 4 weeks p.t., and interestingly, a new developmental stage, the early LiS started to develop from 2 weeks p.t., as evident by the further increase in size (growing wide and stumpy rather than in length) and the formation of the gut, which started out as a bifurcated gut that later fused together in the aboral part of the schistosomula’s body. By week 4 p.t., the late LiS developed, characterized by clearly visible ventral and oral suckers and an elongation of the aboral part of the parasite’s body. Its activity could be either restricted to the oral or aboral part of the body or could encompass the entire worm. This stage was characterized by a steady increase in length (1289.6 ± 247.0 µm) after 6 weeks of culture (S3 Fig). The time point of this development was the same for HM and DMEM. Even though the percentage of the early LiS in DMEM (29.4% or 34.0 ± 2.8 out of 115.5 ± 6.4 total) was higher than that in HM (6.1% or 9.3 ± 1.2 out of 153.0 ± 20.0), the percentage of the late LiS was comparable between DMEM (11.5 ± 5.0 out of 115.5 ± 6.4 total) and HM (15.3 ± 3.2 out of 153.0 ± 20.0) with around 10% having developed after 8 weeks of incubation (Fig 2C, E). Importantly, the growth of the NTS in HM supplemented with HSe was faster compared with that in DMEM since LiS were remarkably bigger (S2 Fig, arrows). Strikingly, despite the addition of HSe, a large number of NTS died in DMEM (S2 Fig, arrow heads). Since in HM, NTS had a higher survival rate, a slightly faster growth rate as well as a yield of 10% LiS juvenile worms after 8 weeks of culture, we decided to use HM for all future experiments.

**Fig 2.**
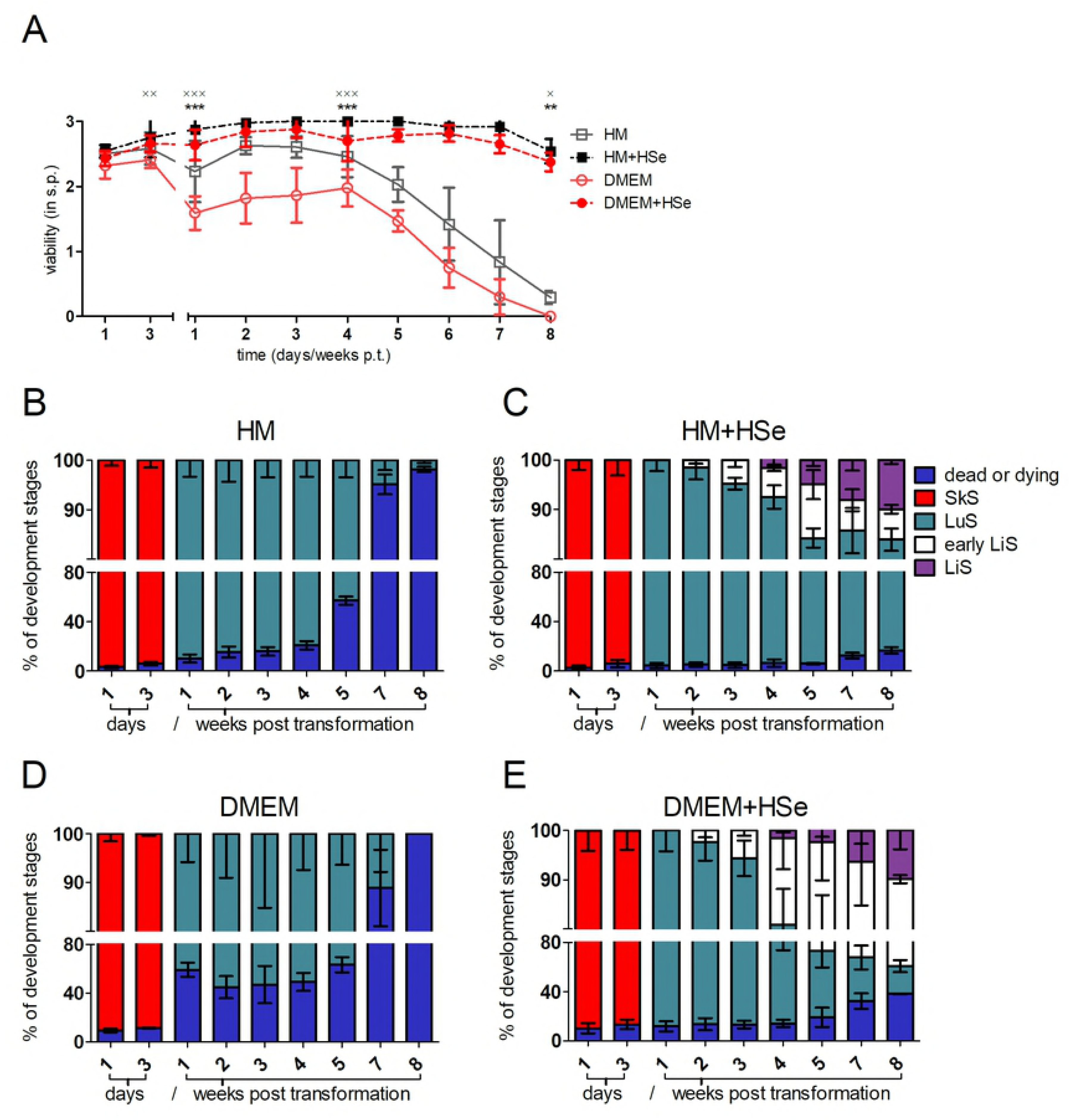
Presence of human serum *in vitro* enhances NTS viability and induces juvenile worm development. NTS were cultured in HM and DMEM with and without 20% HSe. (A) Viability scoring was performed at the indicated time points. The percentage of developmental stages in culture with either HM (B), HM + 20% HSe (C), DMEM (D) or DMEM + 20% HSe was calculated for the indicated time points using worm-stage counts obtained during bright field microscopy. Results are representative of at least three independent experiments. Each data point is shown as a mean ± SD of pooled data from three independent experiments with three biological replicates each. * *p* ≤ 0.05, ** p ≤ 0.01, *** p ≤ 0.001 comparing HM with HM + 20% HSe, × p ≤ 0.05, ×× ≤ 0.01, ××× ≤ 0.001 comparing DMEM with DMEM + 20%. HSe. HSe, human serum; SkS, skin stage; LuS, lung stage; LiS, liver stage; s.p., score points; p.t., post transformation.

### Human serum increases the efficiency of juvenile worm generation in a concentration-dependent manner

Next, we investigated whether HSe induced development of advanced larval stages of *S. mansoni* increases in a concentration-dependent manner. Therefore, we cultured NTS in HM supplemented with 1, 5, 10, 20 or 50% of HSe. Human serum, even at the lowest concentration (1%) used, successfully prevented the death of NTS otherwise starting in the 5^th^ week p.t. (Fig 3A) and survival rates remained steadily above 67% (49.0 ± 7.4 dead in 151.7 ± 15.3 total) throughout the experiments (8 weeks) in all HSe-supplemented conditions (Fig 3B-D). Surprisingly, however, at a concentration of 50% HSe, the survival rate (69.1% or 44.7 ± 10.1 dead out of 144.7 ± 21.2 total) as well as viability (2.2 ± 0.6) started to decline around week 8 (Fig 3D, E). Nevertheless, the final viability score and survival rate was still higher than in the control (viability score of 0.3 ± 0.0 and survival rate of 1.8% or 160.3 ± 7.8 dead out of 163.3 ± 8.5 total) (Fig 3A, E). However, development of NTS up to early and late LiS was only observed at higher concentrations (single early LiS starting in 5% HSe and late in 20% HSe) of HSe (Fig 3C, D and S4 Fig). The early LiS could be observed starting 2 weeks p.t. in all concentrations that allowed its development. However, the number of early LiS increased together with the concentrations of serum. The first parasites in the late LiS were detected at 4 weeks p.t. in 20% HSe and single worms even one week earlier in 50% HSe. The overall percentage of late LiS parasites increased with rising serum concentrations, and, finally, around 10% LiS had developed (15.3 ± 3.2 in 153.0 ± 20.0 total parasite count/well) in 20% HSe compared to approx. 18% (25.3 ± 4.2 in 144.7 ± 21.2 total) in 50% HSe-supplemented HM (Fig 3C, D). Since viability decreased in 50% HSe-supplemented medium after the 7^th^ week p.t. and sufficient numbers (13-19 late LiS/well) of late LiS juvenile worms were generated at 20% HSe [22, 24], we decided to use 20% HSe supplementation for the generation of juvenile worms for drug screening experiments.

**Fig 3.**
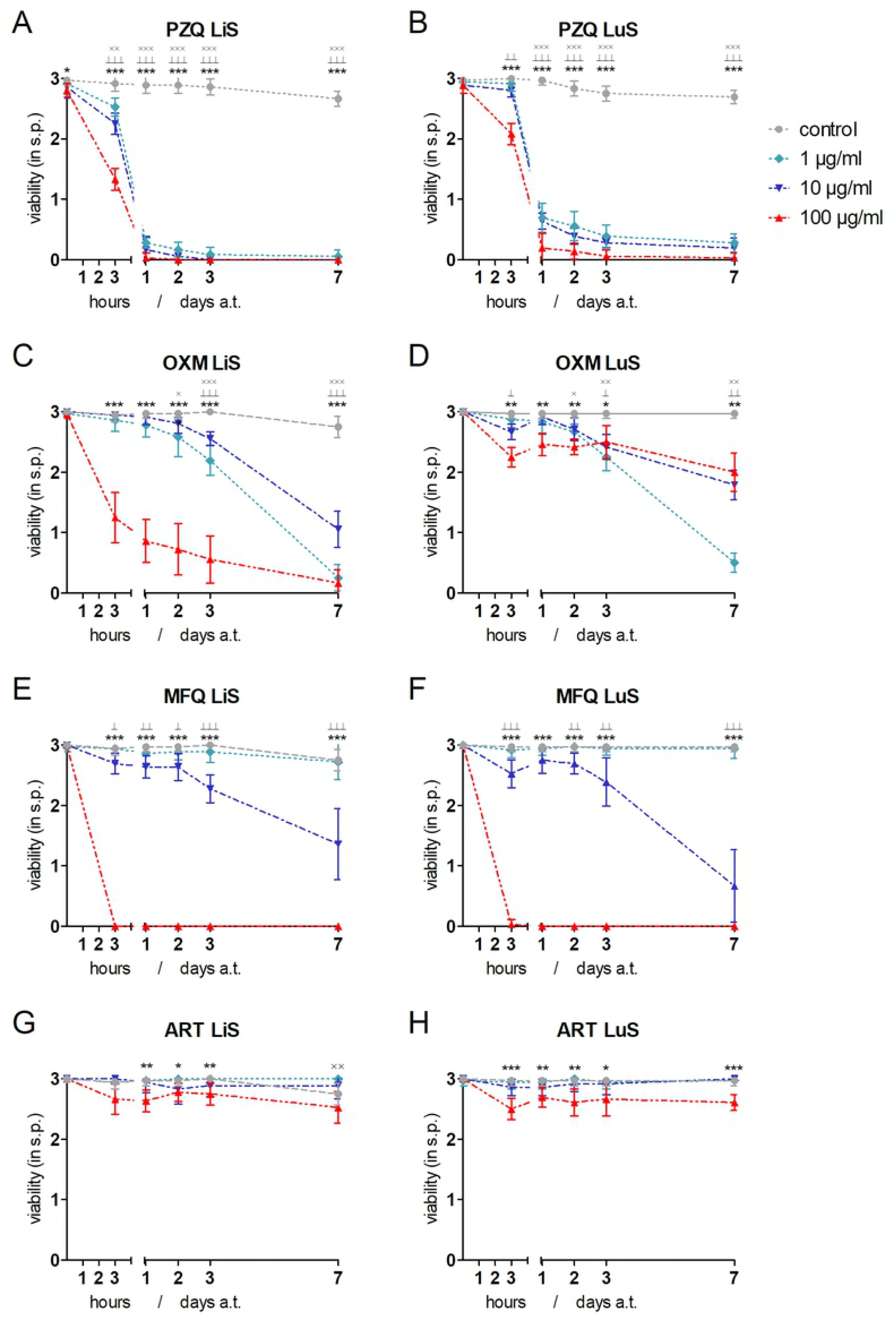
Efficiency of juvenile worm generation in medium supplemented with human serum increases concentration-dependently. NTS were cultured in HM supplemented with 1%, 5%, 10%, 20% and 50% of HSe. The percentages of the developmental stages as well as dead parasites in culture with (A) HM alone or supplemented with (B) 1%, (C) 20% and (D) 50% HSe were calculated at indicated time points and (E) their viability was scored during bright field microscopy. Results are pooled from three independent experiments with at least three biological replicates each. ×× p ≤ 0.01 comparing HM vs. 1% HSe; ^⊥⊥^ p ≤ 0.01 comparing HM vs. 5% HSe; ˅˅ p ≤ 0.01 comparing HM vs. 10% HSe; ≠≠ p ≤ 0.01 comparing HM vs. 20% HSe; ** comparing p ≤ 0.01 HM vs. 50% HSe. HSe, human serum; s.p., score points; p.t., post transformation; SkS, skin stage; LuS, lung stage; LiS, liver stage

### Drugs with known anti-schistosomal properties killed schistosomulae in a stage-dependent manner

Schistosomes have a complex, multi-stage lifecycle. Even within mammalian hosts, their maturation encompasses well-defined stages. We then were interested to test the different stages (SkS, LuS and late LiS) in an *in vitro* drug testing; we chose drugs with known anti-schistosomal properties [24, 28]. We cultured *in vitro* late LiS (6-week-old schistosomulae) (Fig 4A, C, E and G), LuS (7-day-old NTS) (Fig 4B, D, F and H) and SkS (24-hour-old NTS) (S5 Fig A-D) NTS in the presence or absence of different concentrations (100, 10 and 1 µg/ml) of PZQ (Fig 4A, B and S5 Fig A) OXM (Fig 4C, D and S5 Fig B), MFQ (Fig 4E, F and S5 Fig C) or ART (Fig 4G, H and S5 Fig D) followed by regular viability scorings.

**Fig 4.**
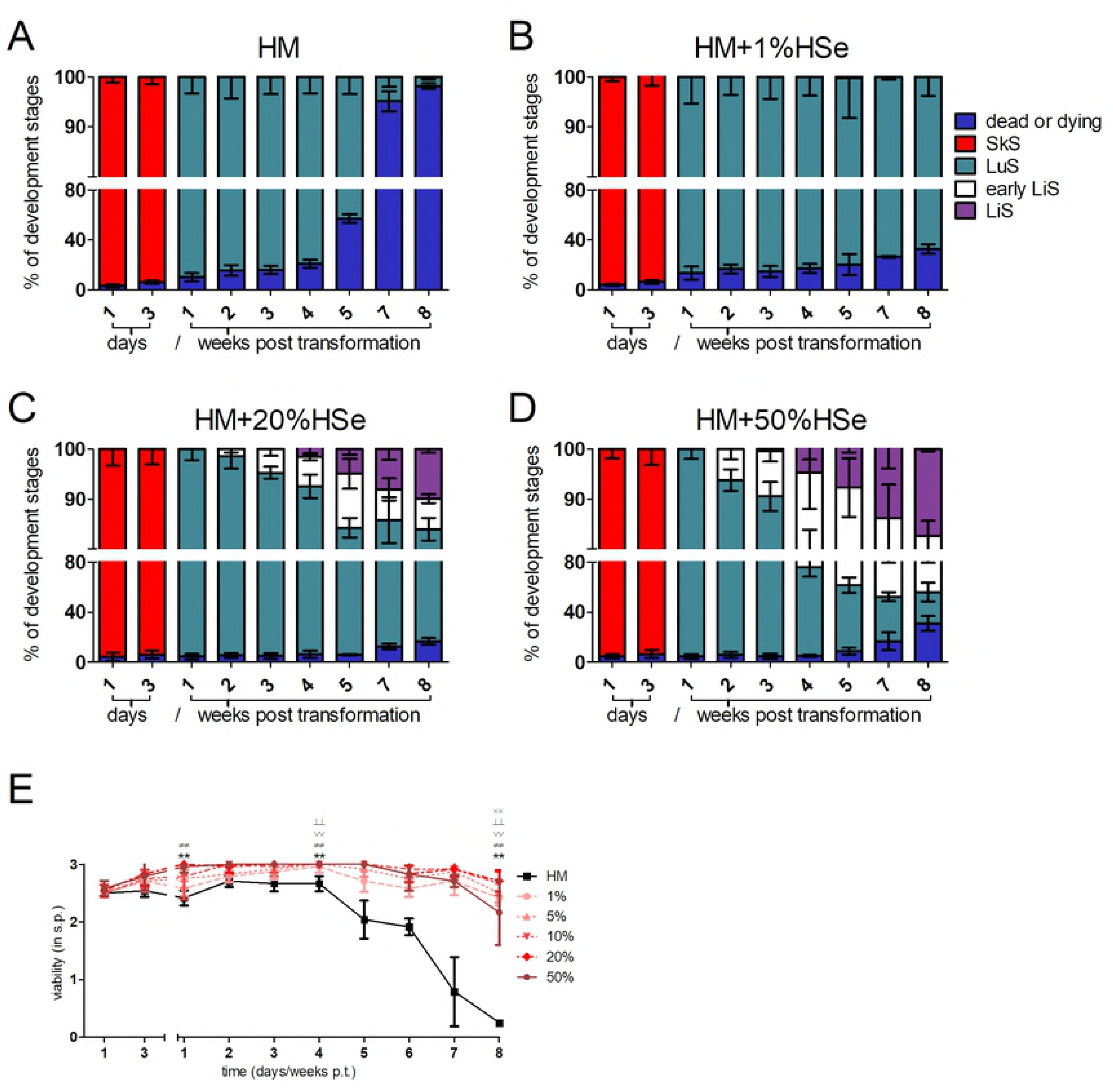
Efficient drug screening of advanced larval stages of *S. mansoni* generated in human serum. NTS were cultured in HM supplemented with 20% HSe (E-H) for 1 week to generate LuS and (A-D) 6 weeks to obtain LiS schistosomulae. (A, E) Praziquantel (PZQ), (B, F) Oxamniquine (OXM), (C, G) Mefloquine (MFQ) and (D, H) Artemether (ART) were added at indicated concentrations for the entire duration of the experiment. 1% DMSO in HM served as control. Viability was scored before treatment (0 h) and 3 hours, 1, 2, 3 and 7 days a.t. Each data point is shown as a mean ± SD of three independent experiments with at least three biological replicates each. (××× p ≤ 0.001, ×× p ≤ 0.01, × p ≤ 0.05 comparing control with 1µg/ml drug; ⊥⊥⊥ p ≤ 0.001 ⊥⊥ p ≤ 0.01 ⊥ p ≤ 0.05 control with 10 µg/ml drug; \*\*\**p* ≤ 0.001, \*\**p* ≤ 0.01 *p ≤ 0.05 control with 100 µg/ml drug). LuS, lung stage; LiS, liver stage; s.p., score points; a.t., after treatment.

PZQ treatment was detrimental to LiS and LuS at all concentrations, which was clearly evident from the drop in viability as soon as 3h a.t. and the death of almost all parasites on day 1 a.t. (Fig 4A, B), characterized by contracted parasites, damaged tegument and no detectable motility (S6 Fig). No such pronounced effect of PZQ was observed on SkS schistosomulae, however, only high concentrations of PZQ slightly reduced the observed viability starting at 3h a.t., but NTS viability then stabilized for the remaining part of the experiment (S5 Fig A). In serum-free conditions, PZQ activity was comparable to serum-supplemented HM as observed on day 3 a.t. At that time point, the susceptibility of the LiS and LuS to PZQ compared with that of the SkS was clearly evident. Serum-free conditions did not alter the activity profile of PZQ on the SkS or LuS (Fig 5A). OXM and MFQ both had similar effects on the LiS in high concentrations. In particular, MFQ had a clear and strong anti-schistosomal activity at 100 µg/ml. In contrast to MFQ, OXM was also potent at 1 µg/ml. even more so than at 10 µg/ml (Fig 4C, E). MFQ showed similar activity on the SkS and LuS compared with the LiS (Fig 4E, F and S5 Fig C). Overall, OXM was potent in reducing the viability of the parasites in the SkS at all tested concentrations. Even though the LuS was the least susceptible stage to OXM, a reduction in the viability could still be observed which was most prominent at 1 µg/ml (Fig 4C, D and S5 Fig B). The stage-dependent vulnerability of the schistosomulae was observed quite well on day 3 a.t. It is worth mentioning that LuS and SkS NTS were slightly more susceptible to OXM in serum-free culture in the concentration of 100 µg/ml. MFQ exerted an increased activity against both SkS and LuS NTS in serum-free compared to HSe-supplemented culture (Fig 5B, C). Closer and careful observations revealed that the highest concentration of OXM induced a hyperactive state directly following the addition of the drug at all stages, which was followed by heavy granulation and tegumental damage in the SkS and LiS already starting on day 1 a.t. and, in the LuS, slightly delayed starting one week a.t. Morphologically, MFQ caused almost instantaneous contraction as well as heavy granulation with a blurred, disintegrated outline of the parasite at 100 µg/ml (S6 Fig). ART did not show any effect on viability or a morphological alteration in LiS or LuS (Fig 4G, H), but a slight reduction in viability of the SkS at 100 µg/ml (S5 Fig D). On day 3 a.t., we could not detect a drop in viability or any morphological changes in any of the tested stages in HSe-supplemented medium (S6 Fig) or in serum-free medium; however, we could detect a clearly schistosomicidal effect at 100µg/ml, with the death following an initial paralysis of the parasites. However, upon addition of the drug to serum free-culture, the previously solved drug precipitated to a small extent (Fig 5D). Taken together, we could show the varying stage-dependent activity profiles of selected compounds already known for their anti-schistosomal properties.

**Fig 5.**
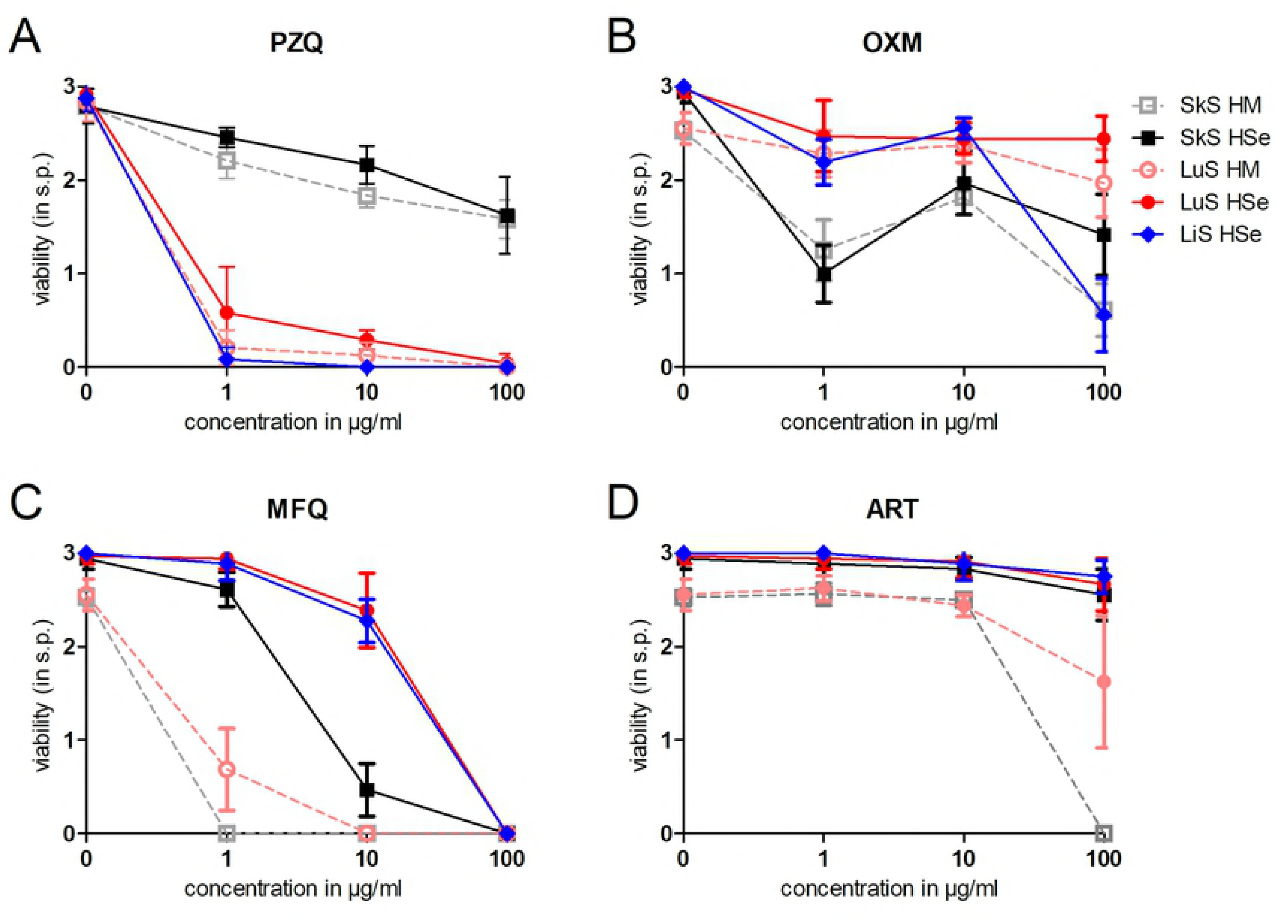
Drug sensitivity is dependent on the developmental stage of *S. mansoni* larvae generated *in vitro*. NTS were cultured and matured in HM alone or supplemented with 20% HSe. 100, 10 and 1 µg/ml of PZQ (A), OXA (B), MFQ (C) and ART (D) were then added to the culture 1 day p.t. to test the SkS, 7 days p.t. to test the LuS and 6 weeks p.t. to test the LiS. Viability was scored 72h a.t. (××× p ≤ 0.001, ×× p ≤ 0.01, × p ≤ 0.05 comparing control with 1 µg/ml drug; ⊥⊥⊥ p ≤ 0.001, ⊥⊥ p ≤ 0.01, ⊥ p ≤ 0.05 comparing control with 10 µg/ml drug; \*\*\**p* ≤ 0.001, \*\**p* ≤ 0.01 *p ≤ 0.05 control with 100 µg/ml drug). SkS, skin stage; LuS, lung stage; LiS, liver stage; PZQ, praziquantel; OXM, oxamniquine; MFQ, mefloquine; ART, artemether; s.p., score points.

## Discussion

In the London Declaration of 2012, the world, represented by partners in governments, academia, NGOs, pharmaceutical companies and more, committed itself to accelerating the control, elimination and eradication of 10 NTDs by 2020 [29]. Despite successes in several areas, progress towards the elimination of schistosomiasis has remained rather limited [30]. Reasons for this lack of progress range from challenges in schistosome vector control to limited developments in drug discovery programs. Rather than developing new compounds, the limited resources are focused towards drug repurposing. The limited interest of pharmaceutical companies in researching entirely new compounds is due to the resource- and time-consuming process to get an approval by the health authorities reviewed in 2014 by Panic et al. [31]. Another limitation in *in vitro* screening used for testing compound libraries is the availability of *in vitro*-generated advanced stage parasites. Therefore, current *in vitro* compound screenings of *S. mansoni* still mostly focus on SkS schistosomulae observed for up to 72 h after treatment for initial hit identification or on adult worms retrieved from infected mice for hit confirmation [14, 32, 33]. This focus leaves a “blind spot” for drug efficacy toward intermediate developmental stages of the parasite. The value of monitoring drug efficacy for initial hit identification at all consecutive stages of parasite development in parallel is clearly illustrated by the narrow, highly stage-specific efficacy of PZQ, which shows effects predominantly against juvenile and adult worms, but still represents the most effective and widely used schistosomicidal drug available [34]. Studies on *ex vivo* harvested LuS schistosomulae sometimes manage to bridge this gap, but are challenging and require a living host (invoking practical as well as ethical aspects) [35].

*In vitro S. mansoni* culture systems obviously can circumvent some of these challenges and allow for the continual observation of consecutive larval stages. Established culture protocols rely on the presence of serum (mainly FCS) for early larval stages [36, 37] and the addition of human blood cells to generate advanced larval and juvenile worm stages [16, 38]. Viability scoring relies on visual assessment of the larvae in culture by bright field microscopy, which still is the gold standard for drug efficacy tests [23, 24]. While the method has its merits, and can be highly effective for dedicated drug efficacy testing, it is very labor intensive. At the same time, microphotographic-based automated analysis [39] using algorithms is complicated by the presence of large numbers of RBCs and/or PBMCs that overlay or colocalize with larvae, making reliable assessment of tegument damage, for example, something for the “trained eye” only. Moreover, the repeated addition of fresh cells from human donors is a factor that is difficult or impossible to fully standardize.

In this study, we generated a novel cell-free *in vitro* assay that allows the development and long-term observation of *S. mansoni* larvae. Mechanically transformed NTS [21, 24] were successfully cultured in a serum- and cell-free culture medium for up to four weeks, allowing for their development to LuS schistosomulae but not further. Supplementation with HSe to the culture broke the developmental block at the LuS and propelled development to juvenile worms, and survival for at least eight weeks. Thus, this cell-free culture for advanced stage *S. mansoni* represents important progress in existing tools that rely on the addition of “non-standardized” human blood cells [16]. Instead, the HM, normally used to cultivate hybridomas and other cell lines, is highly standardized, supplemented with transferrin, insulin, bovine serum albumin (BSA)/oleic acid complex, absorbable amino acids, D-glucose, vitamins and minerals. The presence of albumin and insulin may be critical for the prolonged survival (≥ 4 weeks) of parasites under serum-free conditions. Especially since earlier findings showed that schistosomula can ingest and digest albumin and IgG *in vitro*, utilizing them as nutrient sources [40] and additionally, it was shown that *S. mansoni* has two insulin receptors (SmIR1 and SmIR2), essential for the survival of the pathogen *in vitro* as well as *in vivo* [41]. Interestingly, and in contrast to the natural development *in vivo*, NTS development halted in serum-free media at a stage that, in terms of size and morphology, closely resembles that found in the lung vasculature [17]. Considering that the parasite is blood-dwelling, permanently surrounded by the blood components of its typical definitive human host [2, 27, 42], we were interested to investigate if supplementation with HSe promoted continued NTS development. Indeed, not only did we observe improved overall viability, but also the serum induced concentration-dependent development of NTS to LiS parasites.

Schistosomulae could be maintained for extended periods of time (more than a year) through addition of HSe alone in contrast to non-supplemented controls. However, concentration-dependent advances in stage development (in 50% HSe) came at the price of increased larval mortality at seven weeks and onwards. This may be explained by the increased numbers of parasites reaching the LiS, their increased size and corresponding increased rate in medium nutrient depletion. This would be in line with the fact that the parasite’s feeding and cell divisions are limited, both indicators of low metabolism rates until it reaches the LiS about 15 days after infection *in vivo* [17]. We found supplementation with 20% HSe to provide an optimum between HSe consumption and frequency of medium changes, generating a reasonable number of juvenile worms for further studies. This allows the incorporation of all developmental stages in initial hit identification strategies for future drug screenings, making it possible to identify compounds with an activity profile predominantly targeting juvenile or adult worms as is the case for PZQ [43]. Such drugs would probably otherwise be missed. Taken together, we show that in contrast to cercariae that are susceptible to the complement system present in serum [44], schistosomulae not only tolerate the presence of HSe, but actually require it to progress to late-stage larval stages within their definite host, the human.

Another aim of this study was to investigate the suitability of this method to screen compound libraries for schistosomicidal activity against, in particular, advanced stages of *S. mansoni* in addition to the already often tested early stages [24, 39]. NTS showed comparable stage-specific susceptibility to four drugs known to possess anti-schistosomal properties as seen previously with other *in vitro* assays [16, 24], confirming the validity of our novel culture method. For example, PZQ, undoubtedly the most important tool to control schistosomiasis in the field, [ 16, 45] was described to act predominantly on juvenile and adult worms [19, 43], something we could confirm in our novel culture for the corresponding developmental stages of *S. mansoni* in accordance with previous work [24]. The substantial tegumental damage and anti-schistosomal activity in our assay induced by OXM, the drug used against *S. mansoni* before the discovery of PZQ [46, 47], was also comparable to that observed by others [16, 24]. Specifically, morphological changes of the tegument as well as schistosomicidal activity more pronounced against LiS and SkS schistosomulae at high concentrations could be observed. The observed increased activity in 1 µg/ml compared with 10 µg/ml surprised us. However, similar observations have been made in previous studies [24]. MFQ is an anti-malarial drug, that was shown to also be active against *S. mansoni in vivo* and *in vitro* [48] and is thought to impact heme detoxification as well as glycolysis in the parasite [49]. Interestingly, the stage-dependent activity profile of MFQ is different to that of PZQ and more potently targets the early larval stages than mature worms [25], something we were able to confirm in our novel assay as well. ART is another anti-malarial drug that was shown to act against *S. mansoni* as well [28], thought to act via toxication by the hemin byproduct of the parasite’s digestion of hemoglobin. Indeed, in absence of a cellular source of hemoglobin, ART seemed to be inactive, supporting the notion that hemin is required for the efficacy of the drug [50]. In the absence of serum during early developmental stages, we found strongly enhanced schistosomicidal activity of MFQ, slightly increased potency of ART, but mostly unchanged activity of PZQ and OXM. Bioavailability of MFQ is known to be strongly dependent on binding to plasma proteins (which can be as high as 98% [51]) and for ART plasma-binding lies between 92-98%, in line with our observed increase in activity in serum-free cultures [52]. On the other hand, PZQ binds to a lesser extent to plasma proteins (~80%) in concentrations of 10-100 µg/ml and *in vitro* as low as 50% [53], something that is also reflected in the limited increase in toxicity we observed in non-serum supplemented cultures. Importantly, our assay allows researchers to continually observe toxicity effects and inhibition of maturation from early developmental stages towards mature larval and juvenile worms in settings ranging from compound or drug testing, but also to screen for natural factors from non-permissive hosts or, reversely, identify growth-promoting compounds from the host-adapted, parasite friendly environment.

Taken together, juvenile worms generated in our cell-free assay behave and respond similarly to *ex vivo* harvested worms from infected laboratory animals in drug screening assays [24]. However, in contrast to the limited numbers of labor-intensive, *in vivo* generated parasites, our *in vitro* culture method allows for the generation of good numbers of parasites at all stages of development for large-scale drug screening assays. Also, the independency of host blood cells facilitates automated assessment of larval viability in large-scale assays, due to a lack of visual interference by host cells. In addition, the high level of standardization and minimization of host-to-host variability in the assay, will allow researchers to use it to investigate and identify components within HSe that are exploited by the parasite for its development in the dominant definite human host and define mechanisms that underlie the host-specificity of this parasite. Ultimately, such understanding will pave the way for the identification of new drug and vaccination targets.

## Acknowledgements

We would like to thank Prof. Ulm for advice concerning the statistical analysis of the data, Nermina Vejzagic, PhD for thorough reading and helpful advice concerning the manuscript and Laura Hunt for careful proof-reading of the manuscript.

## Supplements

**S1 Fig. Detailed description of morphological parameters to determine the viability score points of NTS *in vitro***.

Photomicrographs represent indicated viability score points, which results from the overall assessment of motility, morphology and granularity of all NTS per well. Photomicrographs were taken on day 7 post transformation using a digital camera fitted to an inverted microscope (10x).

**S2 Fig. HM enhances survival and accelerates growth compared to DMEM.**

NTS were cultured either in DMEM or HM supplemented with or without 20% HSe. Photomicrographs were taken on day 35 post transformation with a 10x magnification. Arrowheads indicate dead NTS. Arrows indicate LiS (early LiS (DMEM) and late liver stage (HM)).

**S3 Fig. *In vitro* developmental stages from skin stage schistosomula to juvenile worms.**

NTS were cultured in Hybrido Med (HM) supplemented with 20% human serum (HSe). Photomicrographs were taken at indicated time points. For skin, lung and day 21 liver stage, a 10x magnification was used and for the day 35 liver stage, a 5x magnification.

**S4 Fig. Human serum ensures long-term survival of *S. mansoni* larvae already at low concentrations.**

NTS were transformed as described before and kept in HM alone or supplemented with 1%, 5%, 10%, 20% and 50% of human serum. Photomicrographs were taken with a 10x magnification on day 35 post transformation. Arrowheads indicate dead NTS. Arrows indicate LiS (early LiS (5% and 10%) and late liver stage (20% and 50%)).

**S5 Fig. *In vitro* drug testing on skin stage NTS.**

NTS were cultured in HM supplemented with 20% HSe. PZQ (A), OXM (B), MFQ (C) and ART (D) were added at 100, 10 and 1 µg/ml 24 h p.t. and the viability was scored at indicated time points. Statistical analysis was performed by Mann-Whitney U testing comparing drug concentrations to DMSO controls followed by a Bonferroni correction (p.t., post transformation; a.t., after treatment). For statistical analysis, a Mann-Whitney U test was performed comparing drug concentrations to the DMSO control (s.p., score points; a.t., after treatment) (××× p≤0.001, ×× p≤0.01, × p≤0.05 ⊥⊥⊥ comparing control with 1 µg/ml drug; p≤0.001 ⊥⊥ p≤0.01 ⊥ p≤0.05 control with 10 µg/ml drug; \*\*\**p* < 0.001, \*\**p* < 0.01 *p≤0.05 control with 100 µg/ml drug).

**S6 Fig. Drug-induced morphologic changes in all developmental stages.**

NTS were cultured in HM supplemented with 20% HSe. PZQ, OXA, MFQ and ART were added for a final concentration of 100 µg/ml to the culture, HM supplemented with 1% DMSO served as control. Photomicrographs were taken 24h after drug treatment at a 10x magnification.

